# Cisplatin drives mitochondrial dysregulation in sensory hair cells

**DOI:** 10.1101/2024.01.29.577846

**Authors:** David S. Lee, Angela Schrader, Jiaoxia Zou, Wee Han Ang, Mark Warchol, Lavinia Sheets

## Abstract

Cisplatin is a chemotherapy drug that causes permanent hearing loss by injuring cochlear hair cells. The mechanisms that initiate injury are not fully understood, but mitochondria have emerged as potential mediators of hair cell cytotoxicity. Using *in vivo* live imaging of hair cells in the zebrafish lateral-line organ expressing a genetically encoded indicator of cumulative mitochondrial activity, we first demonstrate that greater redox history increases susceptibility to cisplatin. Next, we conducted time-lapse imaging to understand dynamic changes in mitochondrial homeostasis and observe elevated mitochondrial and cytosolic calcium that surge prior to hair cell death. Furthermore, using a localized probe that fluoresces in the presence of cisplatin, we show that cisplatin directly accumulates in hair cell mitochondria, and this accumulation occurs before mitochondrial dysregulation and apoptosis. Our findings provide evidence that cisplatin directly targets hair cell mitochondria and support that the mitochondria are integral to cisplatin cytotoxicity in hair cells.

## Introduction

Cisplatin is a commonly used and effective chemotherapy for the management of solid tumors, but also causes irreversible hearing loss in more than 50% of treated adults and children ^1–3^. This hearing loss decreases quality of life and has additional long-term ramifications in children for its effect on speech and language development ^4–6^. Cisplatin has been used clinically since the 1970s and numerous candidate therapies have been developed in an effort to reduce or eliminate cisplatin ototoxicity ^7^. However, only one therapy has been approved by the United States Food and Drug Administration to mitigate cisplatin-induced hearing loss and its use is limited to children with localized non-metastatic solid tumors; otoprotective drug candidates for adults remain investigational ^8^. Drug discovery has been challenged, in part, by an incomplete understanding of the cellular mechanisms that drive cisplatin-induced hearing loss. Therefore, elucidating the pathophysiology of cisplatin ototoxicity may have substantial clinical implications.

Cisplatin causes damage to cochlear hair cells, the sensory receptors that facilitate hearing, and may do so through a variety of mechanisms. The most studied pathway of cisplatin cytotoxicity is the formation of bulky adducts with nuclear DNA, thereby preventing replication in rapidly dividing cells, such as cancer cells ^9–11^. Yet, hair cells are mitotically inactive, which suggests that another process may drive their vulnerability to cisplatin. More recently, production of pathologic reactive oxygen species (ROS) has been shown to facilitate cisplatin ototoxicity, suggesting that free radical scavengers might mitigate this process ^12,13^. Although multiple pathways may lead to ROS overproduction, mitochondria have emerged as likely mediators of cisplatin ototoxicity because they are a major source of physiologic ROS and represent the only other organelle in hair cells to contain DNA ^14,15^. In support of this theory, preclinical studies show that cisplatin leads to elevated mitochondrial ROS production, increased ratios of Bax to Bcl-2 (mediators of apoptosis and hair cell survival, respectively), and activated caspase-3 in hair cells ^16–18^.

The pattern of cochlear injury in response to cisplatin further implicates mitochondria as critical players in cisplatin ototoxicity. The cochlea is tonotopically organized such that hair cells located at its base transmit high frequencies and those present at its apex transmit low frequencies ^19^. Cisplatin causes hearing loss by initially damaging hair cells at the basal turn of the cochlea followed by progressive hair cell injury towards the apical end ^1,17,20–22^. While the mechanisms that underlie this selective hair cell susceptibility are unknown, tonotopic differences in hair cell metabolic activity, endogenous levels of antioxidants, and intracellular calcium homeostasis may contribute to cisplatin’s distinct pattern of cochlear injury ^20,23,24^. Mitochondria, the organelle responsible for many of these functions, may therefore serve as direct or indirect targets of cisplatin. Similar relationships between mitochondria and high-frequency hearing loss have also been observed in other disease processes, including presbycusis (age-related hearing loss) and aminoglycoside ototoxicity ^25–28^. Altogether, the literature suggests that mitochondrial dysregulation is potentially a key cellular event that drives cisplatin ototoxicity.

Despite an association to ototoxicity, direct observation of cisplatin-induced mitochondrial dysregulation has been elusive. The mammalian cochlea is generally not suitable for live imaging because it is encased in bone and optically inaccessible. In contrast, zebrafish contain hair cells that reside along their surface in a sensory organ called the lateral line. These hair cells are bundled into innervated structures called neuromasts and they are genetically, morphologically, and functionally similar to mammalian hair cells ^29,30^. Since zebrafish larvae are transparent, their lateral-line organ permits consistent drug exposure protocols and live imaging of intact hair cells. These features make zebrafish a powerful model for exploring mechanisms of cisplatin ototoxicity and targeted drug discovery ^31,32^. In the present study, we use transgenic zebrafish with genetically encoded biosensors to explore mitochondrial contributions to selective hair cell susceptibility and to characterize the progression of mitochondrial dysregulation after exposure to cisplatin. Pre- and post-exposure imaging demonstrate that greater cumulative mitochondrial activity increases hair cell vulnerability to cisplatin. Time-lapsed imaging of zebrafish treated with cisplatin demonstrates a slow rise in hair cell mitochondrial calcium and cytosolic calcium followed by a rapid elevation immediately prior to death. We further show that cisplatin accumulates in the mitochondria, and that greater amounts of mitochondrial accumulation of cisplatin is associated with hair cell death. Lastly, we demonstrate that cisplatin accumulation within mitochondria occurs prior to mitochondrial dysregulation and caspase-3 activation. Cumulatively, these results not only provide further evidence that mitochondria are involved in cisplatin ototoxicity, but also suggest that they may be direct targets of cisplatin itself.

## Results

### Greater cumulative mitochondrial activity in hair cells increases susceptibility to cisplatin

As stated above, the mammalian cochlea demonstrates selective injury of hair cells in high frequency regions following cisplatin exposure. We have previously observed that the zebrafish lateral-line organ also experiences selective hair cell death in response to cisplatin, i.e., exposure to cisplatin appears to target a subset of neuromast hair cells in a dose-dependent manner ^33^. Since the lateral-line organ does not exhibit tonotopic organization, selective hair cell injury observed within zebrafish is surprising, especially when considering neuromast hair cells diffusely take up cisplatin within 15 minutes of exposure ^34^. Our initial experiments investigate whether mitochondria play a role in differential hair cell vulnerability to cisplatin.

A prior zebrafish study investigating lateral-line injury after exposure to neomycin, an aminoglycoside antibiotic that also causes high-frequency hearing loss, indicates that cumulative mitochondrial activity contributes to hair cell vulnerability ^23^. To determine whether the lateral-line organ demonstrates a similar pattern of selective susceptibility in response to cisplatin, we performed experiments on transgenic zebrafish expressing the biosensor mitoTimer (*Tg*(*myo6b:mitoTimer*)) in their hair cells ^23^. MitoTimer encodes a DsRed mutant (DsRed1-E5) localized within mitochondria that irreversibly changes from green to red over time as the Tyr-67 residue undergoes dehydrogenation ^35^. Thus, mitoTimer serves as an indicator of cumulative mitochondrial oxidation based on the ratio of red to green fluorescence within individual hair cells ^23,36^. We measured baseline mitoTimer fluorescence in 6 days post fertilization (dpf) transgenic larvae (Figure 1A), then exposed them to 1 mM cisplatin for 2 hours followed by 2 hours of recovery in embryo media (EM). This protocol produces a lesion with approximately 60% hair cell death ^33^. Upon completion of the cisplatin exposure protocol, we obtained post-treatment images and classified each hair cell as alive or dead based on fragmentation and clearance from the neuromast ^23,37^. Baseline red:green ratios of mitoTimer fluorescence were calculated for each hair cell. We observed a significant difference in the median mitoTimer fluorescence ratios between living and dying cells (Living cells: 0.79, interquartile range = 0.48 to 1.00, n = 113 hair cells; dying cells: 1.26, interquartile range = 1.07 to 1.47, n = 73 hair cells; Mann-Whitney U test, p < 0.0001; Figure 1B-D). This result suggests that greater cumulative mitochondrial activity increases vulnerability to cisplatin-induced hair cell death.

**Figure 1.**
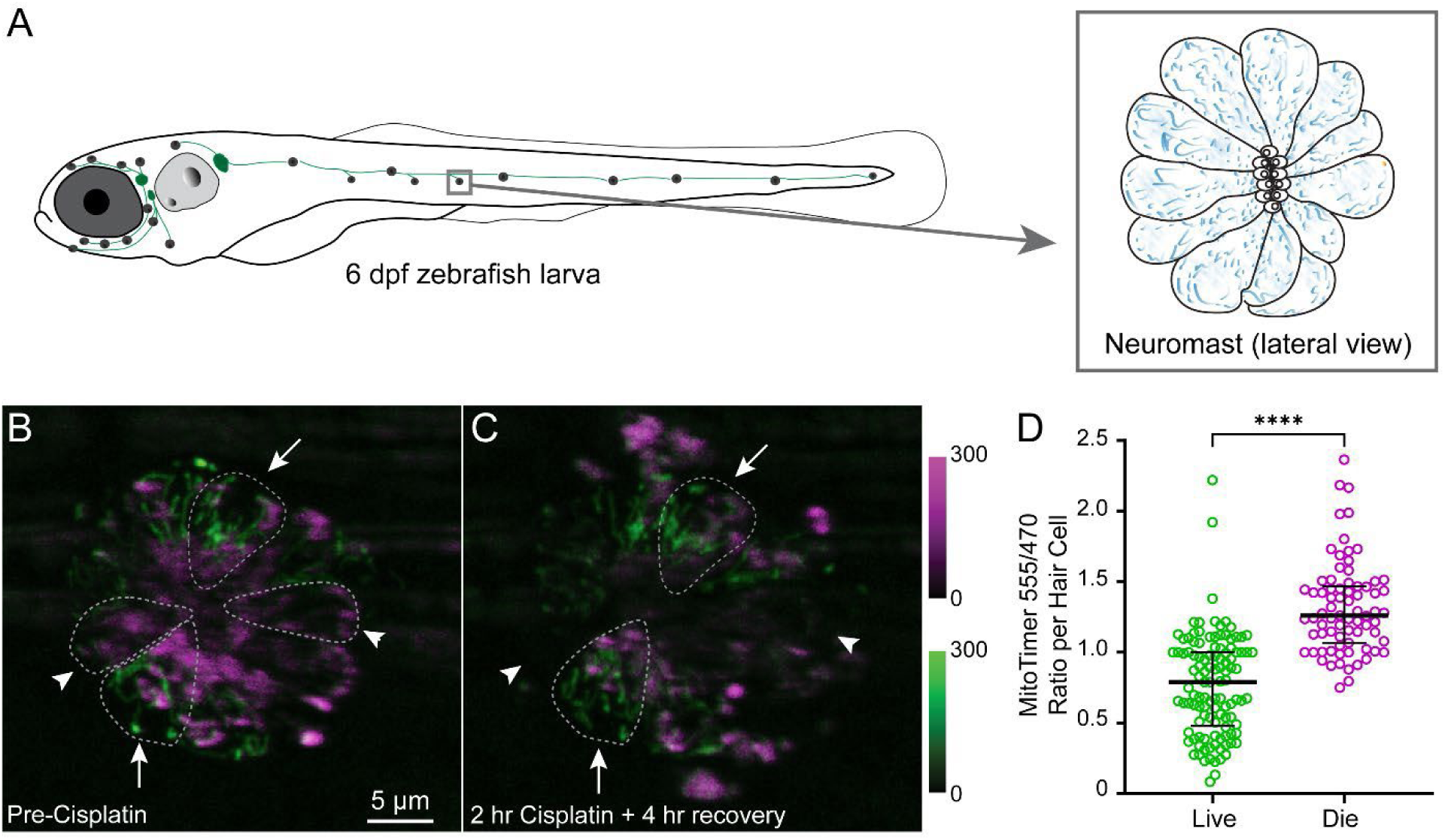
Cumulative mitochondrial activity increases hair cell susceptibility to cisplatin. **(A)** Schematic of larval zebrafish. Lateral line afferent nerves are shown in green, neuromasts are gray spots. Inset shows lateral view of neuromast hair cells with mitochondria shown in blue. **(B – C)** Maximum-intensity projections of confocal z-stack images of *Tg(myo6b:mitoTimer)* neuromasts prior to (**B**) and after (**C**) 2 h cisplatin (1 mM) exposure followed by 2 h recovery in EM. Representative hair cells with high baseline cumulative mitochondrial activity (high 555/470 ratio) are indicated by arrowheads and representative hair cells with low cumulative mitochondrial activity (low 555/470 ratio) are indicated by arrows. **(D)** Baseline ratios of mitoTimer fluorescence indicate that dying hair cells exhibit significantly higher red to green ratios than surviving hair cells (Mann-Whitney U test, \*\*\*\**p* < 0.0001). Median values with corresponding interquartile ranges are shown. n = 113 alive hair cells, 75 dead hair cells; N = 3 trials, 2 zebrafish per trial, 3 neuromasts per zebrafish. Panels (B) and (C) have the same scale bar.

To verify this observation, we sought to determine whether elevating baseline mitochondrial activity prior to cisplatin exposure affects hair cell vulnerability. Therefore, we exposed 5-day-old *Tg*(*myo6b:mitoTimer*) larvae to either continuous water flow in a microflume apparatus or maintained larvae in stationary EM for 24 hours (Figure 2A). We then measured red:green ratios of mitoTimer fluorescence in individual hair cells. Larvae exposed to sustained water current demonstrated increased red:green ratios of mitoTimer fluorescence compared to those maintained in EM (flume cells: 3.21, interquartile range = 1.87 to 4.20, n = 466 cells; no flume cells: 2.08, interquartile range = 1.00 to 3.33, n = 458 cells; Mann-Whitney U test, p < 0.0001; Figure 2 B–C, F).

**Figure 2.**
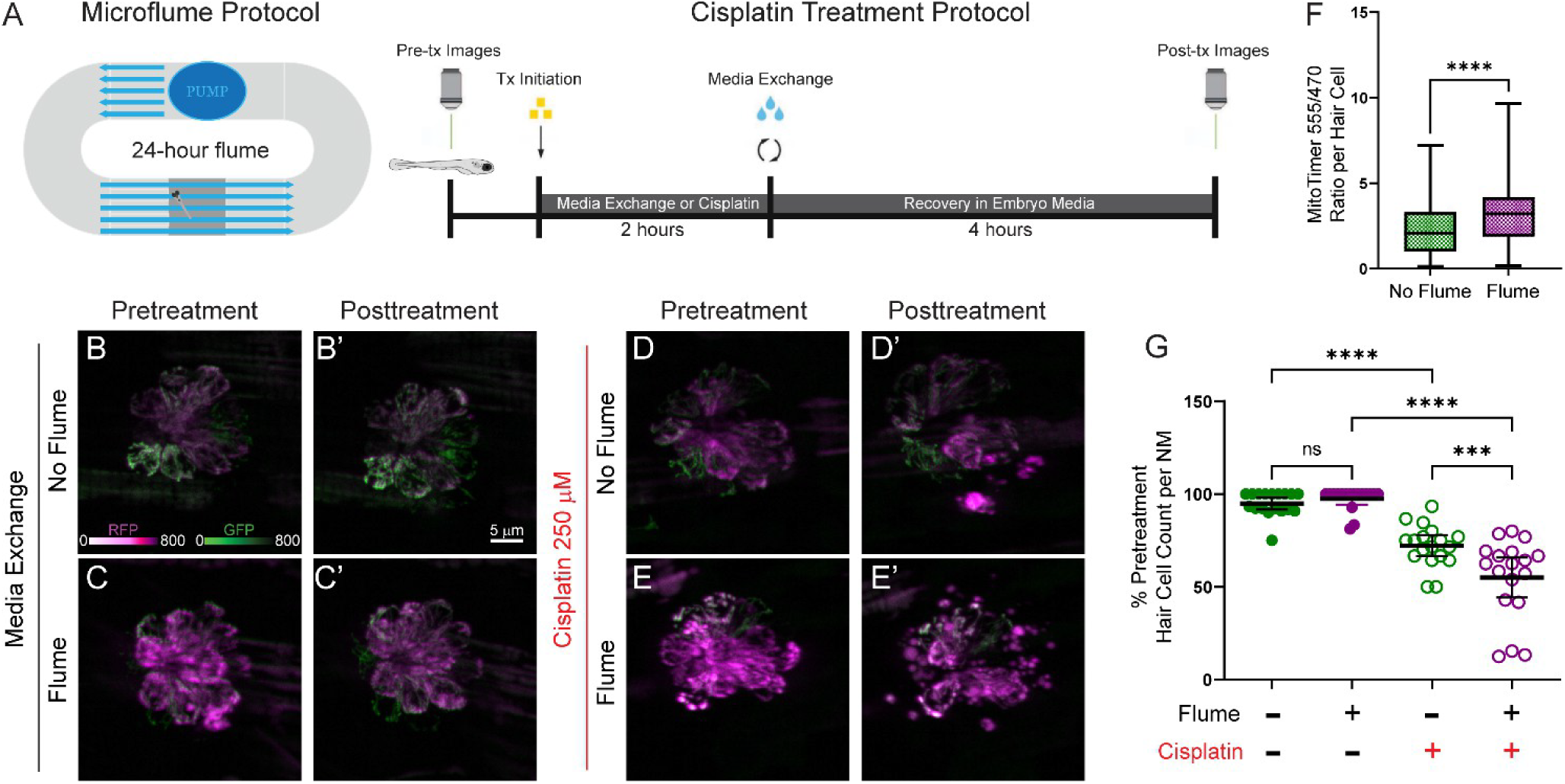
Sustained lateral line hair cell stimulation increases cumulative mitochondrial activity and susceptibility to cisplatin. **(A)** Schematic of the microflume and workflow of the cisplatin treatment protocol. **(B – E)** Maximum-intensity projections of confocal z-stack images of *Tg(myo6b:mitoTimer)* zebrafish neuromasts prior to (**B, C, D, E**) and after 2 h treatment with either EM (media exchange; **B’, C’**) or 250 µM cisplatin (**D’, E’**) followed by recovery in EM for 2 h. Prior to initiating the exposure protocol, zebrafish either underwent incubation with no water flow for 24 h (**B**, **D**) or sustained stimulation to water flow in a flume for 24 h (**C**, **E**). **(F)** Ratios of mitoTimer fluorescence (normalized to no flume condition) after completion of a 24 h incubation in the presence or absence of sustained water flow indicate that prolonged hair cell stimulation increases cumulative mitochondrial activity (Mann-Whitney U test, \*\*\*\**p* < 0.0001). **(G)** Percent of baseline hair cells per neuromast after completion of the cisplatin treatment protocol indicate that moderate-dose cisplatin (250 µM) kills a greater proportion of hair cells among flume stimulated versus unstimulated lateral line neuromasts (Tukey’s multiple comparisons test, adjusted ns = 0.94, ****, *p* < 0.0001; \*\*\**p* = 0.0012). Median values with corresponding interquartile ranges are shown for (F). Mean values with corresponding 95% confidence intervals shown for (**G**). n = 458 hair cells no flume, 466 hair cells in flume; N = 3 trials, 6 zebrafish per condition per trial, 2 – 3 neuromasts per zebrafish.

After establishing that sustained mechanical stimulation increases cumulative lateral-line hair cell mitochondrial activity, we then exposed fish to either media exchange or 250 µM cisplatin for 2 hours followed by 4 hours of recovery in EM (Figure 2A). This protocol produces approximately 30% hair cell death ^33^. A lower concentration of cisplatin was used in these experiments so that relative differences in the percentage of hair cell death observed among flume-exposed fish would be more obvious. Upon conclusion of the cisplatin exposure protocol, we obtained post-treatment images and counted the number of remaining hair cells in each neuromast compared to baseline (Figure 2, B–D, F). While all neuromasts exposed to cisplatin demonstrated a decreased mean percentage of viable hair cells when compared to control neuromasts, those that were also exposed to sustained mechanical stimulation exhibited a significantly lower mean percentage of surviving hair cells relative to neuromasts in fish not exposed to sustained water current (Media exchange-no flume: 96.7%, 95% confidence interval = 91.7% to 98.2%, n = 18 neuromasts; media exchange-flume: 97.5%, 95% confidence interval = 94.4% to 100%, n = 17 neuromasts; cisplatin-no flume: 72.1%, 95% confidence interval = 66.6% to 77.7%, n = 18 neuromasts; cisplatin-flume: 55.1%, 95% confidence interval = 44.3% to 65.8%, n = 18 neuromasts; Tukey’s multiple comparisons test, adjusted ns = 0.94, ***adjusted *p* = 0.0012, ****adjusted *p* < 0.0001; Figure 2G). These results indicate that greater cumulative mitochondrial activity increases hair cell vulnerability to cisplatin.

### Influx of mitochondrial calcium precedes hair cell death following cisplatin exposure

Mitochondria have been implicated in cisplatin cytotoxicity as sources of pathologic ROS and initiators of caspase-3-mediated apoptosis in both cancer and non-cancer models ^16–18,38,39^. However, the cellular events and time course of mitochondrial dysregulation have not been well characterized. Our next set of experiments use live *in vivo* imaging of transgenic zebrafish with genetically encoded biosensors to investigate the time course of cisplatin-induced mitochondrial dysregulation.

Mitochondria play a key role in intracellular calcium homeostasis ^40^. We therefore evaluated the time course of changes in mitochondrial calcium levels in response to cisplatin. These experiments utilized transgenic zebrafish that stably express calcium indicator, GCaMP3, in the mitochondria of their hair cells (*Tg(myo6b:mitoGCaMP3);* ^37^). We exposed transgenic larvae at 5 – 7 days post-fertilization (dpf) to either media exchange or 1 mM cisplatin for 2 hours and then allowed recovery in EM for an additional 2 hours. Baseline mitoGCaMP3 fluorescence was measured before treatment initiation and images were acquired every 10 minutes throughout the duration of the cisplatin exposure protocol (Figure 3A). We then determined the change in mitoGCaMP3 fluorescence relative to baseline (ΔF/Fο) of individual hair cells at each time point. Next, we classified cells as dead or alive based on their pattern of hair cell mitochondrial fragmentation and extrusion. We aligned data to the time at which dying hair cells were cleared from the neuromast (Figure 3B–D). Cisplatin-exposed hair cells demonstrated a slight elevation in mitochondrial calcium levels compared to those that underwent media exchange alone (control). However, dying hair cells exhibited a gradual elevation of mitochondrial calcium that peaked immediately prior to hair cell death (Figure 3D). This phenomenon was not observed in living or control hair cells. When evaluating peak changes in mitochondrial calcium levels, the median change in maximum fluorescence of hair cells was significantly greater among dying hair cells than living or control hair cells (control cells: 1.18, interquartile range = 1.03 to 1.44, n = 233 cells; alive cells: 2.23, interquartile range = 1.58 to 3.33, n = 136 cells; dead cells: 4.14, interquartile range = 2.82 to 5.65, n = 155; Dunn’s multiple comparisons test, ****adjusted *p* < 0.0001; Figure 3E). To evaluate when surges in mitochondrial calcium levels occurred, we measured the time at which half the maximum change in fluorescence was observed (t_1/2_ maximum ΔF/Fο) relative to neuromast clearance. This occurred, on average, 36.3 minutes prior to hair cell fragmentation (95% confidence interval = 32.9 to 39.6 minutes, Table 1). Altogether, these data suggest that cisplatin leads to a slow accumulation of hair cell mitochondrial calcium followed by a rapid elevation immediately prior to death.

**Figure 3.**
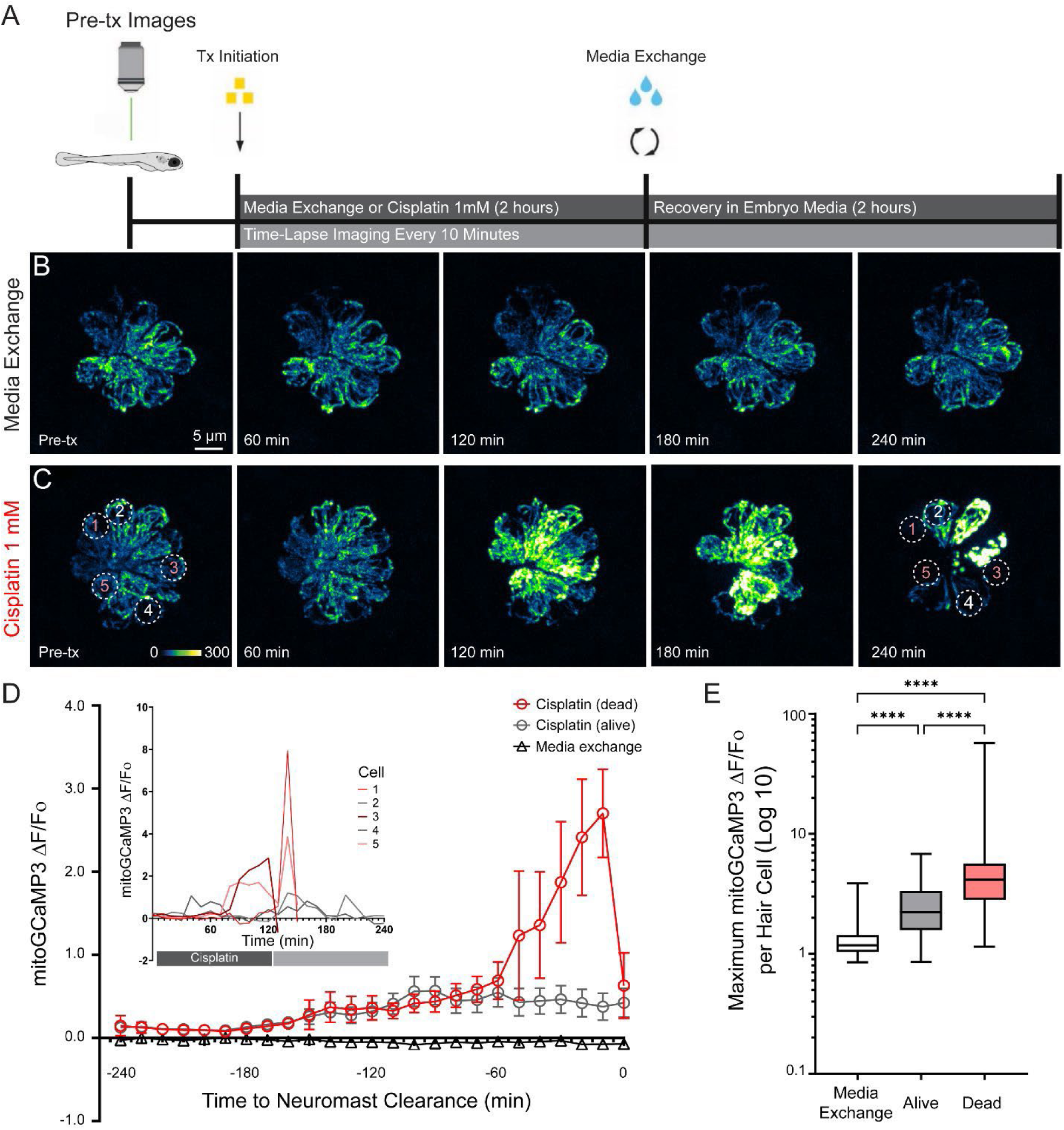
Elevated mitochondrial Ca^2+^ uptake is observed in all cisplatin exposed hair cells and progressively increases prior to hair cell death. **(A)** Schematic of cisplatin exposure and *in vivo* live imaging workflow. **(B – C)** Maximum-intensity projections of time-lapse confocal z-stack images show *Tg(myo6b:mitoGCaMP3)* zebrafish neuromasts at various points throughout the exposure protocol. Zebrafish were treated with either 2 h of EM (media exchange; **B**) or 1 mM cisplatin (**C**) followed by 2 h recovery in embryo media. Dotted circles indicate a representative 4 µm region of interest (ROI) in an individual hair cell used to measure mean mitoGCaMP3 fluorescence intensity. An ROI was applied to every hair cell within each neuromast. **(D)** Changes in mitoGCaMP3 fluorescence intensity relative to mean baseline fluorescence intensity (ΔF/Fο) for each neuromast hair cell over time. Hair cells were grouped by treatment. Cisplatin-exposed hair cells were further categorized into those alive or dead at the conclusion of the exposure protocol. Data are aligned relative to when dying hair cells were cleared from the neuromast. Inset shows changes in mitoGCaMP3 fluorescence intensity in individual hair cells (numbered in C) over the time course of the experiment. **(E)** Maximum change in fluorescence relative to baseline (normalized to control) demonstrates significant differences in mitochondrial Ca^2+^ levels between all three groups of hair cells (Dunn’s multiple comparisons test, ****adjusted *p* < 0.0001). Median values with corresponding interquartile ranges are shown for both (**D**) and (**E**). n = 233 media exchange hair cells, 136 alive cisplatin-exposed hair cells, 155 dead cisplatin-exposed hair cells; N = 3 trials, 3 zebrafish per trial, 2 – 3 neuromasts per zebrafish. **Movie 1.** The corresponding time lapse video of Figure 3C showing changes in hair cell mitochondrial Ca^2+^ during and after cisplatin exposure.

**Table 1.**
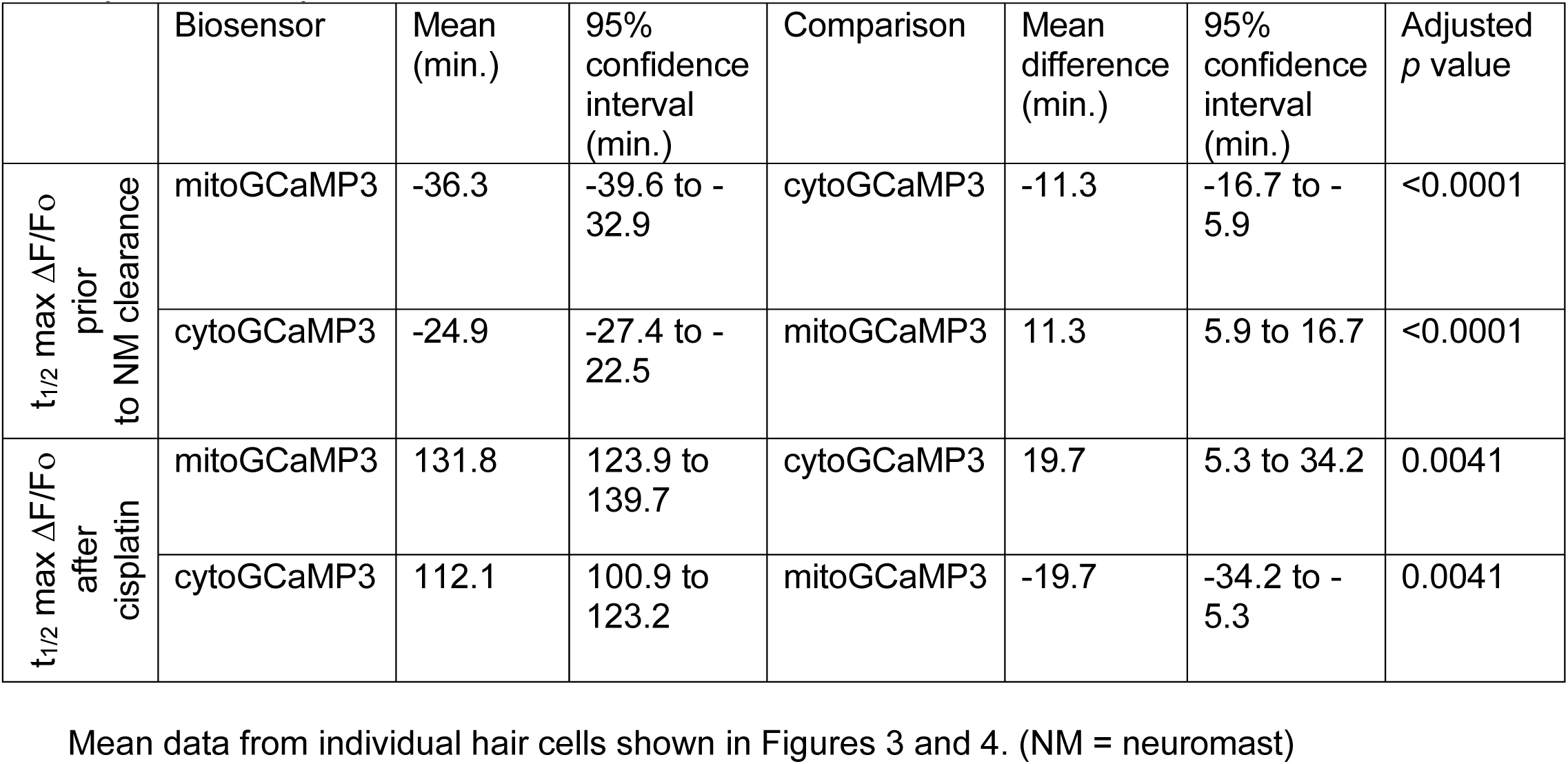
Time to half maximum ΔF/Fο prior to clearance from neuromast and after exposure to cisplatin.

|  | Biosensor | Mean (min.) | 95% confidence interval (min.) | Comparison | Mean difference (min.) | 95% confidence interval (min.) | Adjusted *p* value |
| --- | --- | --- | --- | --- | --- | --- | --- |
| $t_{1/2}$ max $\Delta F/F_0$ prior to NM clearance | mitoGCaMP3 | -36.3 | -39.6 to -32.9 | cytoGCaMP3 | -11.3 | -16.7 to -5.9 | <0.0001 |
|  | cytoGCaMP3 | -24.9 | -27.4 to -22.5 | mitoGCaMP3 | 11.3 | 5.9 to 16.7 | <0.0001 |
| $t_{1/2}$ max $\Delta F/F_0$ after cisplatin | mitoGCaMP3 | 131.8 | 123.9 to 139.7 | cytoGCaMP3 | 19.7 | 5.3 to 34.2 | 0.0041 |
|  | cytoGCaMP3 | 112.1 | 100.9 to 123.2 | mitoGCaMP3 | -19.7 | -34.2 to -5.3 | 0.0041 |

Mean data from individual hair cells shown in Figures 3 and 4. (NM = neuromast)

### Calcium accumulates in cytosol of dying hair cells after exposure to cisplatin

We next explored the time course of cytosolic calcium shifts in response to cisplatin. These experiments utilized transgenic zebrafish that express GCaMP3 in the cytosol of hair cells (*Tg(myo6b:cytoGCaMP3);* ^37^). We measured baseline cytoGCaMP3 fluorescence of transgenic larvae at 5 – 7 dpf, exposed them to either media exchange or 1 mM cisplatin followed by 2 hours of recovery in EM, and obtained time-lapse images at 10-minute intervals (Figure 4A–B). The ΔF/Fο of individual hair cells at each time point was determined and these changes in fluorescence were aligned to time of neuromast clearance (Figure 4C). Among dying hair cells, we observed a slow rise in cytosolic calcium levels followed by a sharp peak immediately before fragmentation. Similar to the observed surge of mitochondrial calcium in dying hair cells (Figure 3), large shifts in cytosolic calcium were not observed in control or living hair cells. The median change in maximum fluorescence over baseline was significantly greater among dying hair cells than living or control hair cells (control cells: 1.12, interquartile range = 1.04 to 1.25, n = 169 cells; alive cells: 1.31, interquartile range = 1.14 to 1.55, n = 155 cells; dead cells: 1.90, interquartile range = 1.36 to 2.41, n = 87 cells; Dunn’s multiple comparisons test, ****adjusted *p* < 0.0001; Figure 4D). The t_1/2_ maximum ΔF/Fο of cytoGCaMP3 occurred 24.9 minutes prior to clearance from the neuromast (95% confidence interval = 22.5 to 27.4 minutes, Table 1). This shift in cytosolic calcium took place 11.3 minutes after the accumulation of mitochondrial calcium (95% confidence interval = 5.9 to 16.7 minutes, Tukey’s multiple comparisons, ****adjusted *p* < 0.0001, Table 1). These data suggest that dying hair cells experience a large increase in cytosolic calcium levels shortly after surges in mitochondrial calcium levels.

**Figure 4.**
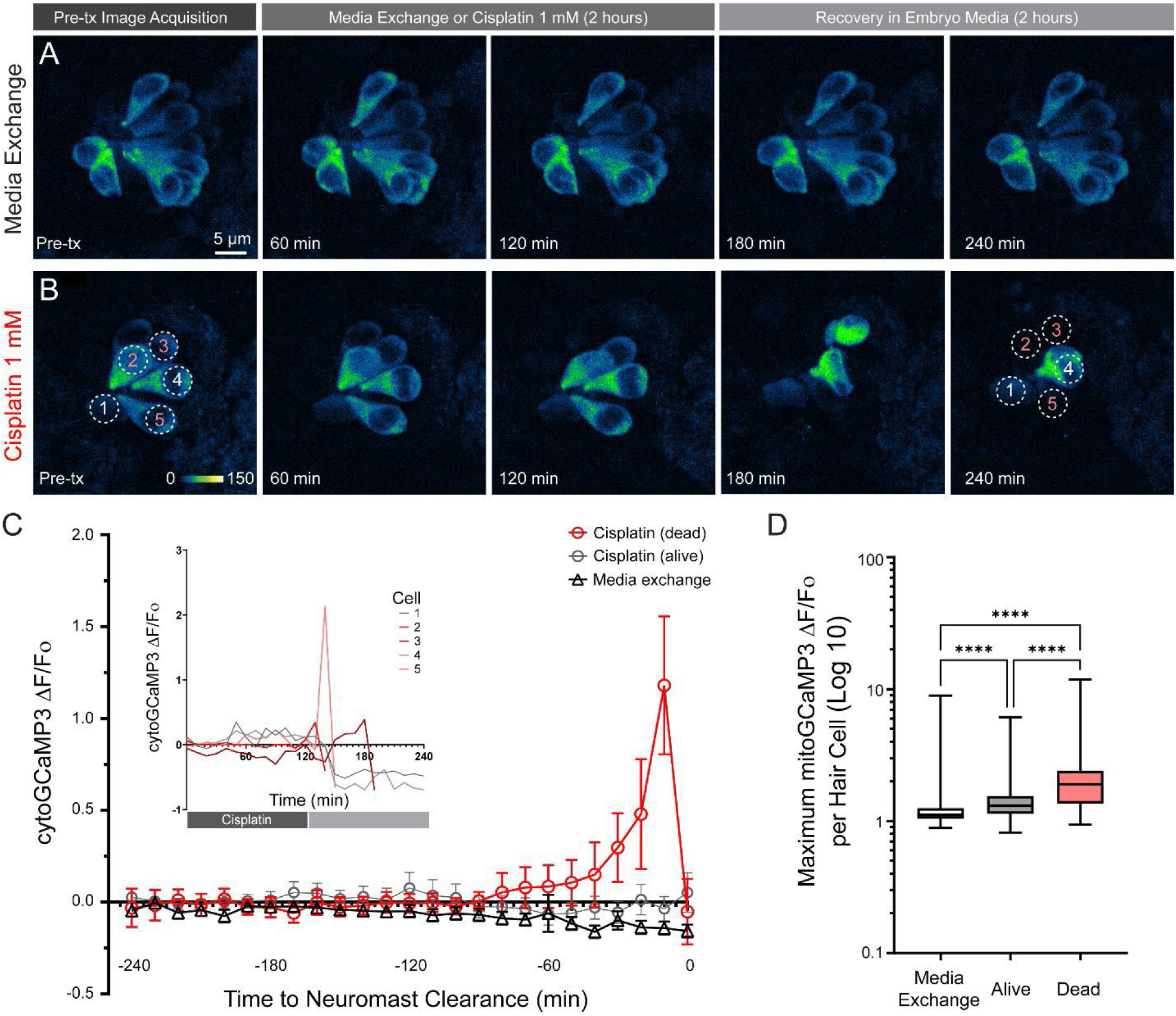
Elevated cytosolic Ca^2+^ uptake precedes cisplatin-induced hair cell death. **(A – B)** Maximum-intensity projections of time-lapsed confocal z-stack images show *Tg(myo6b:cytoGCaMP3)* zebrafish neuromasts at various points throughout the exposure protocol. Zebrafish were treated with either 2 h of EM (media exchange; **A**) or 1 mM cisplatin (**B**) followed by 2 h recovery in EM. Dotted circles indicate representative 4 µm ROI used to measure mean cytoGCaMP3 fluorescence intensity. **(C)** Changes in cytoGCaMP3 fluorescence intensity relative to baseline (ΔF/Fο) for each hair cell over time. Data are aligned relative to when dying hair cells were cleared from the neuromast. Inset shows changes in cytoGCaMP3 fluorescence intensity in individual hair cells (numbered in B) over the time course of the experiment. **(D)** Comparison of maximum change in fluorescence relative to baseline (normalized to media exchange) demonstrates significant differences in cytosolic Ca^2+^ accumulation between all three groups of hair cells (Dunn’s multiple comparisons test, ****adjusted *p* < 0.0001). Median values with corresponding interquartile ranges are shown for both (**C**) and (**D**). n = 169 media exchange hair cells, 155 alive cisplatin-exposed hair cells, 87 dead cisplatin-exposed hair cells; N = 2 trials for media exchange and 4 trials for cisplatin 1 mM, 3 zebrafish per trial, 2 – 3 neuromasts per zebrafish. **Movie 2.** The corresponding time lapse video of Figure 4B showing changes in hair cell cytosolic Ca^2+^ during and after cisplatin exposure.

### Cisplatin treatment leads to progressive activation of caspase-3 prior to hair cell death

We have previously shown that cisplatin induces caspase-3-mediated apoptosis in conjunction with disruption of hair cell mitochondrial bioenergetics, suggesting a mitochondrially driven cell death pathway ^18^. To evaluate the onset and pattern of caspase-3 activation, we exposed wildtype larvae at 6 dpf to either media exchange or 1 mM cisplatin for 2 hours, then allowed recovery in EM for an additional 2 hours. At baseline and every 30-minute interval, a subset of larvae was euthanized, fixed, and stained for activated caspase-3 (Figure 5A–B). We quantified the percentage of hair cells with activated caspase-3 present within a given neuromast at each time point. We then averaged these percentages within each larva to account for fish-to-fish variability in response to cisplatin. Cisplatin-exposed larvae demonstrated significantly greater percentages of hair cells with activated caspase-3 than control fish at approximately 90 minutes after treatment (Figure 5C, mixed model, Bonferroni’s multiple comparisons test, *adjusted *p* = 0.016, **adjusted *p* = 0.0012, ****adjusted *p* < 0.0001; n = 42 – 45 neuromasts per time point from 14 – 15 fish; Table 2). These results indicate that activation of caspase-3-mediated apoptosis begins in a subset of hair cells within 90 minutes of 1 mM cisplatin treatment and peaks at ∼210 minutes after initiating treatment protocol. Cumulatively, these observations suggest activation of the intrinsic apoptosis pathway precedes mitochondrial dysfunction and subsequent hair cell death.

**Figure 5.**
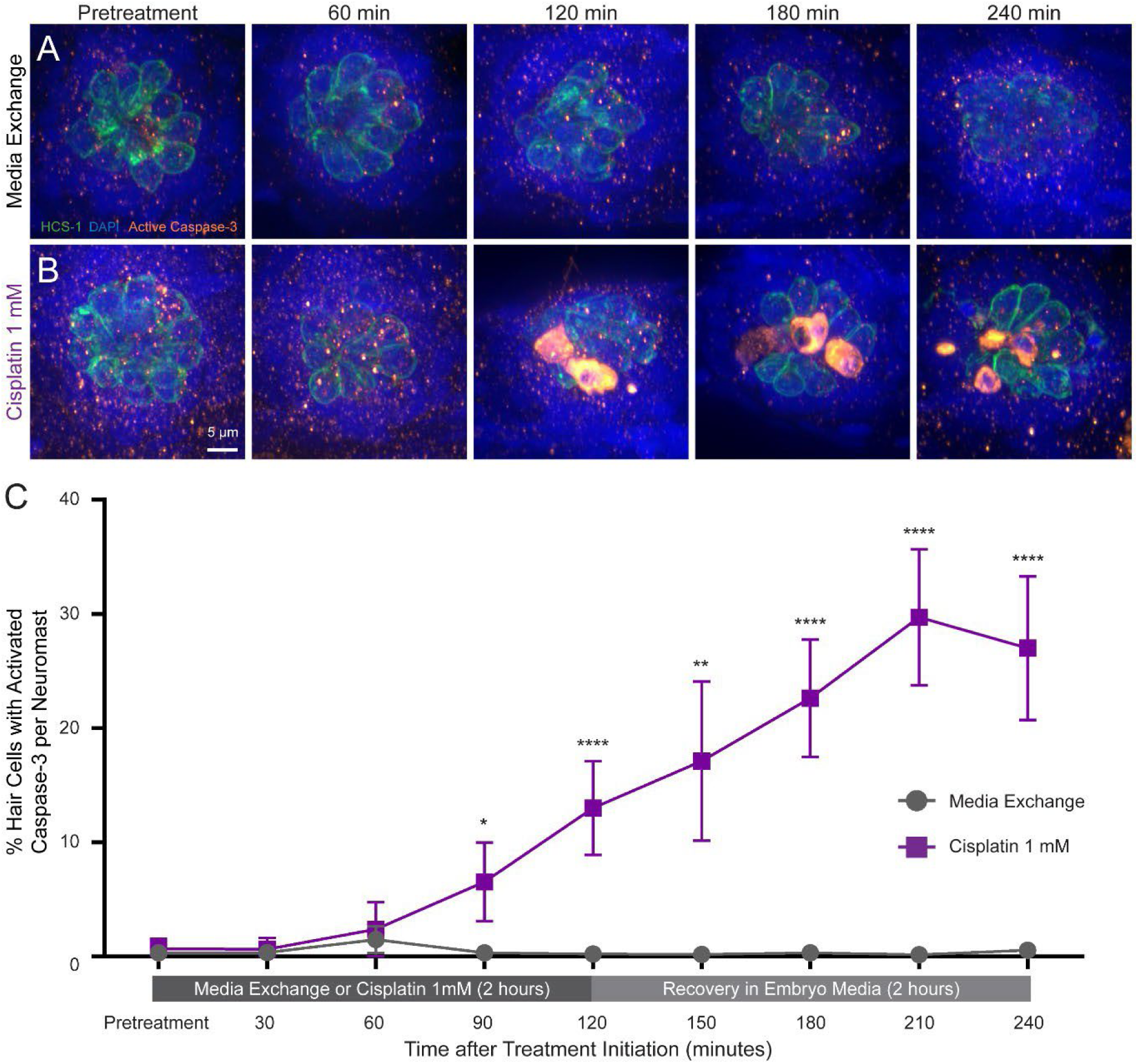
Progressive activation of Caspase-3 in a subset of neuromast hair cells initiates within 90 minutes of 1 mM cisplatin exposure. **(A – B)** Maximum-intensity projections of z-stack confocal images of immunolabeled neuromasts at various points throughout the cisplatin exposure protocol. Zebrafish were treated for various amounts of time with either media exchange (**A**) or 1 mM cisplatin (**B**) followed by recovery in embryo media for up to 2 h. Hair cells were identified by immunolabel of Otoferlin (Hair Cell Soma-1; green) and stained for cleaved Caspase-3 (orange). All cell nuclei were labeled with DAPI (blue). **(C)** Percentage of hair cells per neuromast with activated Caspase-3 throughout the cisplatin exposure protocol. Neuromasts of cisplatin-treated zebrafish demonstrate a significantly greater proportion of hair cells with cleaved Caspase-3 starting at 90 minutes following exposure to cisplatin (Bonferroni’s multiple comparisons test, **adjusted *p* = 0.016, ***adjusted *p* = 0.0012, ****adjusted *p* < 0.0001; Table 2). Mean values with corresponding 95% confidence intervals shown. n = 42 – 45 neuromasts per group; N = 3 trials, 4 – 5 zebrafish per trial, 2 – 3 neuromasts per zebrafish.

**Table 2.** Results from repeated measures mixed model of caspase-3 activation in the presence or absence of cisplatin 1 mM.

| Time after assay initiation (min.) | Exposure group (% HC with activated Caspase-3/NM) |  | Mean difference (%) | 95% confidence interval (%) | Adjusted *p* value |
| --- | --- | --- | --- | --- | --- |
|  | Cisplatin 1 mM | Media exchange |  |  |  |
| 0 | 0.67 | 0.33 | 0.34 | -1.1 to 1.8 | >0.9999 |
| 30 | 0.63 | 0.34 | 0.29 | -1.3 to 1.9 | >0.9999 |
| 60 | 2.4 | 1.48 | 0.9082 | -2.9 to 4.7 | >0.9999 |
| 90 | 6.55 | 0.33 | 6.22 | 0.92 to 11.5 | 0.016 |
| 120 | 13.0 | 0.22 | 12.78 | 6.5 to 19.1 | <0.001 |
| 150 | 17.1 | 0.17 | 16.9 | 6.3 to 27.6 | 0.0012 |
| 180 | 22.6 | 0.32 | 22.3 | 14.4 to 30.2 | <0.001 |
| 210 | 29.7 | 0.16 | 29.5 | 20.5 to 38.6 | <0.001 |
| 240 | 27.0 | 0.56 | 26.5 | 16.8 to 36.1 | <0.001 |

Mean percentages of hair cells with activated-Caspase 3 from neuromasts shown in Figure 5. (HC = hair cell, NM = neuromast)

### Cisplatin heterogeneously accumulates in hair cell mitochondria prior to mitochondrial dysregulation and caspase-3 activation

The prior time-lapse imaging experiments demonstrate that rapid mitochondrial dysfunction occurs immediately prior to hair cell death. This dysfunction may occur due to a variety of reasons, like disrupted endoplasmic reticulum calcium shuttling or ROS overproduction ^25,28,41^. Yet to be determined was whether hair cell mitochondria are direct targets of cisplatin or if mitochondrial dysregulation is purely collateral damage from different cellular processes. Prior literature indicates that cisplatin accumulates in the mitochondria of cancer cell lines, presumably exerting its anticancer effects through adduct formation with mitochondrial DNA ^42^. Yet, accumulation in mitochondria has not been demonstrated within hair cells. We therefore used Rho-Mito, a mitochondria-targeting probe that fluoresces when bound to cisplatin, to explore the uptake of cisplatin in hair cell mitochondria of live zebrafish ^42^. To identify the location of mitochondria, we used the transgenic zebrafish reporter line t*g(myo6b:mitoGCaMP3)* (^37^; see Figure 3). We exposed larvae at 5 – 7 dpf to either media exchange or 1 mM cisplatin for 1 hour, then incubated them in Rho-Mito 3 µM for 15 minutes. Next, baseline images were acquired at this time to observe overlap between Rho-Mito and mitoGCaMP3 fluorescence. The larvae were then allowed to recover for an additional 4 hours in EM. We obtained post-treatment images at the conclusion of the cisplatin exposure protocol and classified each hair cell as alive or dead as previously described ^23,37^. Baseline fluorescence of Rho-Mito was normalized to the median of living hair cells. We first noticed that Rho-Mito fluorescence varies between hair cells, suggesting that mitochondria heterogeneously take up cisplatin. Furthermore, hair cells that died following 4 hours recovery demonstrated a significantly higher median of Rho-Mito fluorescence than hair cells that lived (Dead cells: 14.1, interquartile range = 6.16 to 48.7, n = 92 hair cells; living cells: 5.60, interquartile range = 4.57 to 7.41, n = 229 hair cells; Mann-Whitney U test, p < 0.0001; Figure 6E – F). We also observed heterogeneity in hair cell mitochondrial calcium 1 hour following exposure to cisplatin. However, differences in mitochondrial calcium levels at this timepoint did not correlate with cisplatin ototoxicity (Mann-Whitney U test, ns = 0.14; Figure 6F). This result is consistent with what we reported in Figure 3C, where we observed heterogeneous mitoGCaMP3 fluorescence in individual hair cells at baseline, and minimal changes in mitochondrial calcium following 60 minutes cisplatin exposure. These data reveal that mitochondrial accumulation of cisplatin corresponds with hair cells that subsequently die following cisplatin exposure.

**Figure 6.**
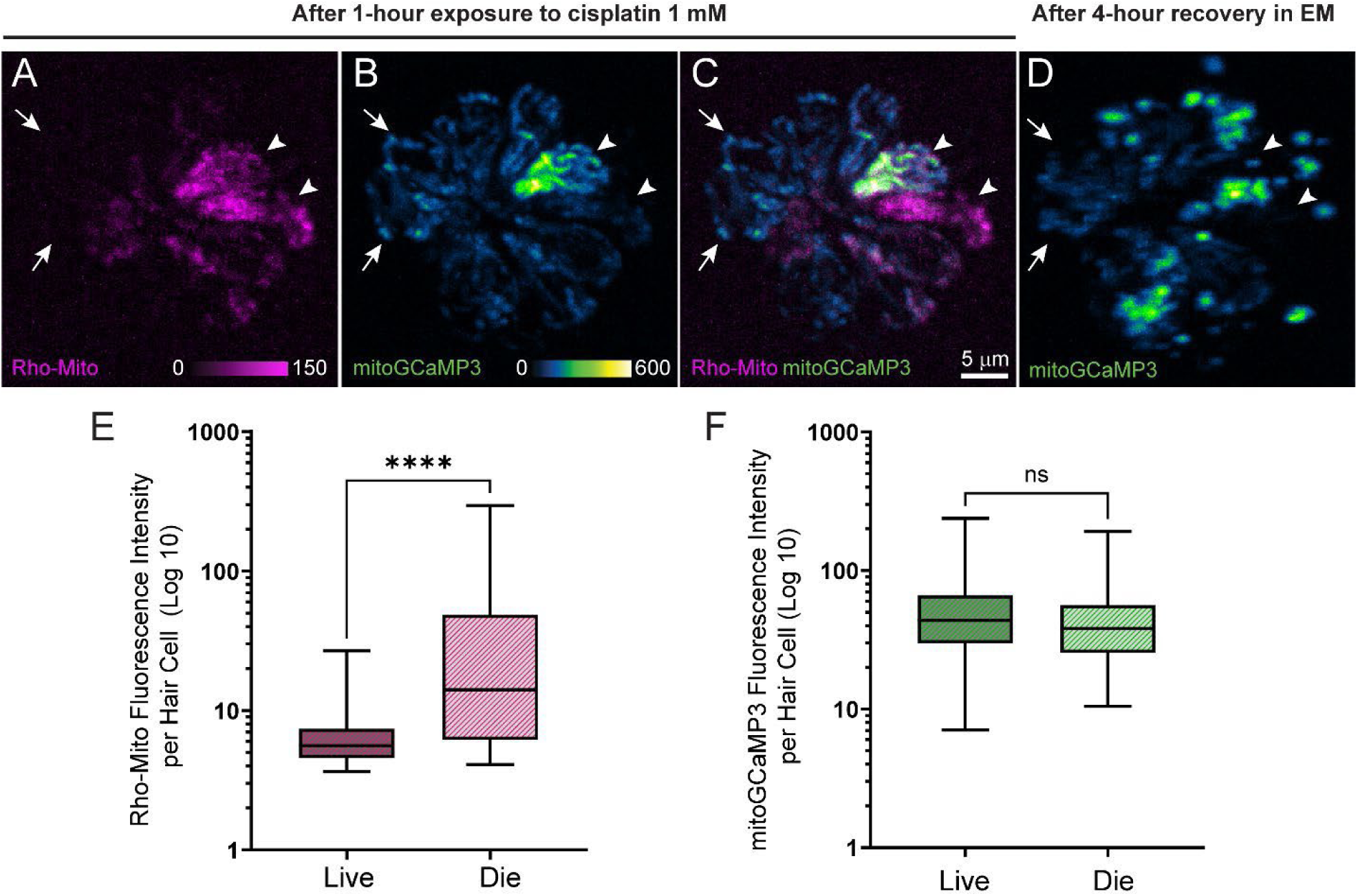
Mitochondrial accumulation of cisplatin corresponds with hair cell death. **(A – D)** Maximum-intensity projections of the same *Tg(myo6b:mitoGCaMP3]* neuromast after exposure to 1mM cisplatin for 1 h (**A – C**) followed by an additional 4 h recovery in EM (**D**). Rho-Mito fluorescence is observed within some, but not all, hair cell mitochondria. Hair cells with elevated Rho-Mito fluorescence following incubation in cisplatin are indicated by arrowheads and hair cells with minimal Rho-Mito fluorescence are indicated by arrows. **(E)** Dying hair cells exhibit significantly more Rho-Mito fluorescence than living hair cells (Mann-Whitney U test, \*\*\*\**p* < 0.0001). (F) MitoGCaMP3 fluorescence in dying hair cells is not significantly different than living hair cells immediately following 1 h cisplatin exposure (Mann-Whitney U test, ns = 0.14). Median values with corresponding interquartile ranges are shown. n = 229 alive hair cells, 92 dead hair cells; N = 3 trials, 2 – 3 zebrafish per trial, 2 – 3 neuromasts per zebrafish. The scale in panels A – D are the same.

A previous study using fluorescently labeled cisplatin suggests diffuse uptake into hair cells within 15 minutes following exposure ^34^. To confirm heterogeneous accumulation of cisplatin in mitochondria and explore the time course of mitochondrial localization, we performed a follow-up experiment in transgenic zebrafish that do not express a reporter in mitochondria to ensure that Rho-Mito localization in mitochondria was not influenced by mitoGCaMP3. We used transgenic fish expressing GFP in neuromast supporting cells to provide a silhouette of hair cells as previously described (*Tg(TNKS1bp1:EGFP);* ^18,43,44^). We exposed 5 – 7 dpf larvae to 1 mM cisplatin for various lengths of time, incubated them in 3 µM Rho-Mito, then acquired images (Figure 7A-C). Neuromast hair cells exposed to cisplatin demonstrated significantly greater median Rho-Mito fluorescence compared to control (Control hair cells: 7.40, interquartile range = 5.61 to 13.41, n = 287 hair cells; 30-minute exposure: 10.68, interquartile range = 6.14 to 29.59, n = 337 hair cells; 60-minute exposure: 13.81, 7.24 to 29.69, n = 320 hair cells; Dunn’s multiple comparisons test, ****adjusted *p* < 0.0001; Figure 7D). Neuromast exposed to cisplatin demonstrated heterogeneous mitochondrial accumulation, with a subset of hair cells accumulating cisplatin within their mitochondria (Figure 7B-C). On post-hoc analysis, hair cells incubated in cisplatin for 60 minutes did not demonstrate increased accumulation of Rho-Mito fluorescence compared to those incubated for only 30 minutes (Dunn’s multiple comparisons test, adjusted ns = 0.26; Figure 7D). Since caspase-3 activation is observed in a subset of hair cells ∼90 minutes after onset of cisplatin treatment (Figure 5), these data indicate that mitochondrial accumulation of cisplatin occurs prior to events that signal hair cell death. Altogether, these data suggest that 1) mitochondrial accumulation of cisplatin occurs in some, but not all, hair cells within the first 30 minutes of exposure, 2) greater accumulation of mitochondrially-localized cisplatin corresponds with hair cell death, and 3) mitochondrial accumulation of cisplatin precedes caspase-3 activation and mitochondrial dysregulation.

**Figure 7.**
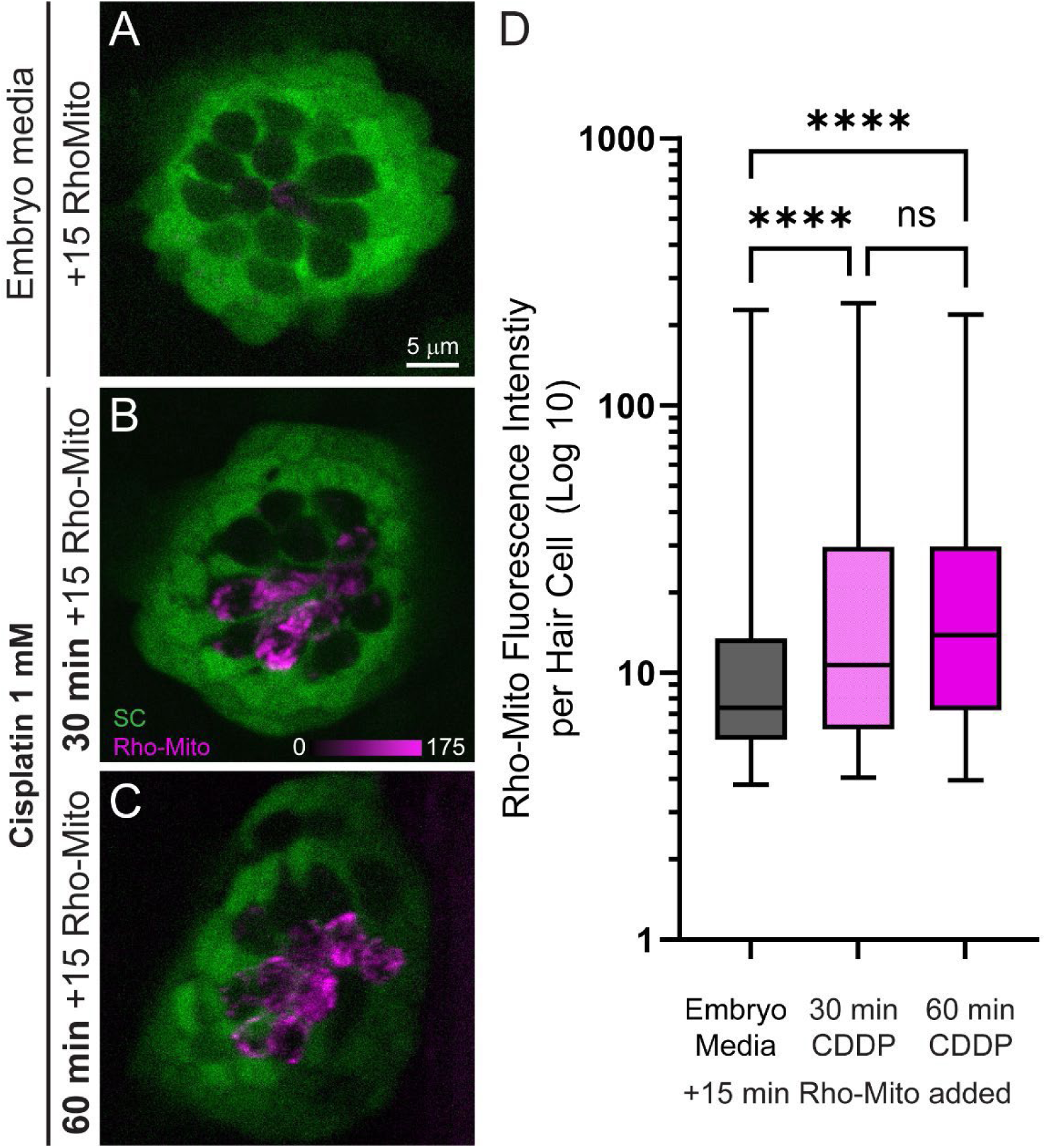
Cisplatin accumulates within hair cell mitochondria shortly after exposure. **(A – C)** Maximum-intensity projections of confocal z-stack images show (*Tg(TNKS1bp1:EGFP)* larvae expressing GFP in neuromast supporting cells at various time points after exposure to cisplatin. Zebrafish were treated with either EM (**A**) or 1 mM cisplatin for up to 1 h (**B** – **C**) followed by a 15 min incubation in 3 µM Rho-Mito. Hair cells were identified as GFP-negative silhouettes. **(D)** Relative fluorescence of Rho-Mito in hair cells normalized to the median of the control. Hair cells exposed to cisplatin demonstrate a significantly greater amount of Rho-Mito fluorescence at all time points (Dunn’s multiple comparisons test, ****adjusted *p* < 0.0001). Although cisplatin appears to progressively accumulate in mitochondria over time, this trend was not significant (Dunn’s multiple comparisons test, adjusted ns > 0.26). Median values with corresponding interquartile ranges are shown. n = 287 media exchange hair cells, 337 hair cells after 30 min cisplatin, 320 hair cells after 60 min cisplatin; N = 3 trials for control and cisplatin-exposed groups, 2 – 3 zebrafish per trial, 2 – 3 neuromasts per zebrafish.

## Discussion

Our recent studies in zebrafish lateral-line hair cells provided evidence that cisplatin disrupts mitochondrial homeostasis and initiates the intrinsic apoptosis pathway through activation of the downstream effector caspase-3 in zebrafish lateral-line hair cells ^18^. In this study, we evaluated whether cumulative mitochondrial activity is associated with hair cell susceptibility to cisplatin, examined the time course of hair cell calcium dysregulation, and determined whether cisplatin accumulates within hair cell mitochondria following exposure. These data reveal that hair cell susceptibility to cisplatin-induced death correlates with both cumulative mitochondrial activity and accumulation of cisplatin within mitochondria, and that mitochondrial accumulation of cisplatin is an early event that precedes activation of caspase-3. Additionally, we show that mitochondrial and cytosolic calcium, which are elevated in all cisplatin treated hair cells, gradually increase then surge prior to hair cell death. Taken together, these results suggest that cisplatin directly targets hair cell mitochondria, and that accumulation of cisplatin within hair cell mitochondria drives mitochondrial dysregulation and contributes to hair cell death.

### Cumulative hair cell mitochondrial activity may underlie selective vulnerability to cisplatin

Our evaluation of cisplatin toxicity in the zebrafish lateral line indicates that hair cells with higher cumulative metabolic activity are more vulnerable to cisplatin-induced death. This observation corresponds with two recent studies evaluating susceptibility of lateral line hair cells to the ototoxic antibiotic neomycin. The first study, which established and validated the hair cell expressing mitoTimer transgenic line used in this study, revealed that hair cells with greater cumulative mitochondrial activity were more susceptible to neomycin-induced death ^23^. A subsequent study reported that hair cells with reduced mitochondrial oxidation and ROS buildup were less vulnerable to neomycin ^45^. Taken together, work in zebrafish lateral-line hair cells supports that cumulative oxidation of mitochondria and accumulation of ROS could underlie the specific susceptibility of hair cells to ototoxic insults.

How does information from zebrafish lateral line studies relate to hair cell susceptibility in humans? In patients that experience ototoxicity from cisplatin treatment, hearing impairment manifests as high frequency hearing loss ^46^. This observed clinical outcome is due to cisplatin preferentially affecting outer hair cells at the basal turn of cochlea, corresponding to the region responsible for detection of high frequency sounds. Preferential ototoxicity to hair cells at the base of the cochlea could be due to unequal distribution of cisplatin within the cochlea, with higher accumulation of chemotherapy at the base than apex ^1,47,48^, and/or intrinsic vulnerability of high frequency outer hair cells to cisplatin. Our results suggest differential hair cell vulnerability to cisplatin is, at least in part, related to differences in mitochondrial metabolism. Observations from cochlear studies in mammalian model systems support this idea. High frequency outer hair cells have been shown to be metabolically more responsive to their microenvironment than low frequency outer hair cells ^49,50^, and genetic disruption of mitochondrial bioenergetic homeostasis makes outer hair cells more susceptible to cisplatin-induced death ^51,52^.

### Hair cell ability to regulate intracellular calcium homeostasis may be predictive of vulnerability to cisplatin

Intracellular calcium homeostasis may contribute to hair cell susceptibility to environmental and ototoxic insults ^53^. Mitochondria play a critical role in regulating local intracellular calcium concentrations as well as provide ATP to drive clearance of cytosolic calcium via plasma membrane Ca^2+^-ATPases (PMCA1 & PMCA2a; ^54^). Sustained clearance of calcium from the stereocilia bundle via PMCA2a pumps is required to maintain calcium at physiological levels ^55^. Hair cells in the high frequency region of the cochlea show larger amplitude current through mechanotransduction (MET) channels corresponding with larger calcium influx. This generates a larger calcium load entering the stereocilia of high frequency hair cells, resulting in greater levels of metabolic stress on mitochondria ^53^. Therefore, the hair cell’s ability to handle intracellular calcium load may be predictive of hair cell susceptibility to environmental and ototoxic stress. Our data shows increased vulnerability to cisplatin with cumulative mitochondrial activity (Figure 2) as well as dysregulation of calcium homeostasis in all hair cells exposed to cisplatin (Figure 3,4), possibly indicating that hair cells with greater cumulative mitochondrial activity have a reduced capacity to regulate intracellular calcium concentrations and are more susceptible to death from dysregulation driven by cisplatin. This idea is further supported by the observation that hair cell mitochondrial calcium measurements at baseline were not predictive of cisplatin vulnerability (i.e. that mitochondria with less cumulative metabolic activity are better able to buffer intracellular calcium levels even when stressed; Figure 6). Inhibiting mitochondrial calcium uptake with either calcium chelators or pharmacological inhibitors has been shown to significantly reduce neomycin induced hair cell death in lateral line neuromasts ^28,37^. Still to be determined is whether modulating mitochondrial calcium transients during a critical window prior to the activation of caspase-3 could provide protection from cisplatin induced hair cell death. One important difference between neomycin-induced and cisplatin-induced hair cell death in the zebrafish lateral line is that neomycin kills hair cells within minutes, while cisplatin-induced death takes hours ^33^. A key challenge to performing this experiment in cisplatin-treated zebrafish is determining when and how to inhibit hair cell mitochondrial calcium uptake such that it provides protection while minimally impacting normal hair cell respiration and metabolism ^56^.

An open question not directly addressed by this study is: what is the link between mitochondrial turnover, mitochondrial calcium levels, and susceptibility to cisplatin-induced hair cell death? Previous data acquired from hair cell expressing mitoTimer fish provides some clues ^23^. This study, which initially generated and characterized the transgenic line, made several important observations with regards to hair cell activity, mitochondrial calcium, and mitochondrial turnover : i) using mitochondrial and cytosolic calcium indicator lines and a 10 Hz pressure wave, they observed mitochondrial calcium levels exhibit a delayed rise time and slow decay in response to stimulation, ii) they reported higher mitoTimer red;green ratios corresponded with MET activity (mutants lacking MET or pharmacological block of MET showed decreased red:green ratios), iii) using the photoconvertible protein mitoEos to track hair cell mitochondrial turnover, they observed less mitochondrial turnover in wildtype fish with intact MET than in fish mutants lacking MET. Cumulatively, these observations suggest that hair cell MET activity contributes to increased mitochondrial calcium, cumulative hair cell oxidation and, surprisingly, reduced mitochondrial turnover. In the context of this previous study, our observations suggest that cumulative mitochondrial stress paired with reduced mitochondrial turnover may contribute to increased hair cell vulnerability to cisplatin-induced injury. This premise is further supported by studies showing faster mitochondrial turnover and increased mitochondrial biogenesis in cancer cells resistant to cisplatin chemotherapy ^57,58^.

### Cisplatin accumulation in mitochondria correlates with hair cell death

We observed mitochondrial accumulation of cisplatin in a subset of hair cells as an early event predictive of hair cell death (Figure 6, 7). This accumulation may trigger downstream cellular events, like surges in mitochondrial calcium, that culminate in hair cell apoptosis. One likely target for cisplatin in hair cells is mitochondrial DNA. Prior work has reported cisplatin-induced mitochondrial DNA damage and mitochondrial degradation in dorsal root ganglion neurons ^59^. Additional studies in head and neck cancer cell lines implicate mitochondrial DNA as a major determinant of cisplatin sensitivity ^60,61^. Moreover, studies evaluating the effects of cisplatin on mitochondria isolated from rat liver suggest that cisplatin directly increases calcium-induced mitochondrial permeability transition and disrupts mitochondrial bioenergetics ^62^, and provide evidence that mitochondrial membrane proteins, such as Voltage-dependent anion-selective channel 1 (VDAC-1), may also be directly targeted by cisplatin ^60^.

However, alternative explanations for mitochondrial dysregulation independent of mitochondrial accumulation of cisplatin exist. Although we expect that hair cells with high Rho-Mito fluorescence will experience a gradual increase in mitochondrial calcium followed by a spike immediately prior to hair cell death, we do not specifically investigate changes in mitochondrial calcium levels among Rho-Mito labeled hair cells. Therefore, processes that cause mitochondrial stress, like overproduction of ROS from NADPH oxidase-3 isoform and xanthine oxidase activation as well as reduction is antioxidant levels, may lead to pathologic shifts in calcium instead of mitochondrial accumulation of cisplatin^22,63–65^. Future work is needed to identify mitochondrial target of cisplatin in sensory hair cells and to determine how accumulation of cisplatin in mitochondria contributes to hair cell death.

## Conclusion

Cisplatin will continue to be a staple for solid and hematologic malignancies in the coming decades. As such, developing strategies that ameliorate or prevent ototoxicity should remain a focus of research and innovation. Using live *in vivo* imaging of zebrafish lateral line organs, this study established that mitochondrial accumulation of cisplatin precedes mitochondrial calcium dysregulation and eventual caspase-3-mediated hair cell death. Our findings advance the notion that mitochondria are key mediators of cisplatin ototoxicity, and that they may even be direct targets of cisplatin itself. Future investigations may seek to determine whether prevention of calcium dysregulation reduces ototoxicity and to characterize the activity of cisplatin within mitochondria. Such studies will provide further insight to the underpinnings of cisplatin ototoxicity and guide additional targeted drug discovery.

## Methods

### Sex as a biological variable

Sex cannot be determined or predicted in larval zebrafish. Larvae were therefore chosen at random without consideration of sex.

### Zebrafish husbandry and lines

Embryos were maintained in embryo media (EM: 15 mM NaCl, 0.5 mM KCl, 1 mM CaCl_2_, 1 mM MgSO_4_, 0.015 mM KH_2_PO_4_, 0.042 mM Na_2_HPO_4_, 0.714 mM NaHCO_3_) at 28°C with 14-hour light and 10-hour dark cycles ^66^. After 3 days post-fertilization (dpf), larvae were transferred to 250-mL plastic beakers, raised in 100 – 200 mL of EM, and fed rotifers daily. All experiments were performed on larvae aged 5 – 7 dpf

Previously described transgenic lines included: w119Tg (*Tg(myo6b:mitoGCaMP3)*; ^28^), w78Tg (*Tg(myo6b:cytoGCaMP3);* ^37^), w208Tg (*Tg(myo6b:mitoTimer);* ^23^, and y229Gt (*Tg(TNKS1bp1:EGFP);* ^43^. The first three lines were used to evaluate changes in mitochondrial calcium levels, cytoplasmic calcium levels, and cumulative mitochondrial activity, respectively. The last line expresses GFP in neuromast supporting cells, permitting identification of hair cells in the absence of genetically encoded biosensors. Fluorescent transgenic larvae were identified at 3 – 4 dpf under anesthesia with 0.01% tricaine in EM.

### Zebrafish live imaging and cisplatin exposure protocol

Larvae at 5 – 7 dpf were anesthetized in 0.01% tricaine methanesulfonate (MS-222) in EM and mounted lateral-side up in an open bath image chamber (Warner Instruments; RC-26G) with a coverglass bottom. A slice anchor (Warner Instruments; 64-0253) was used to secure up to 3 larvae at a time. Larvae were maintained in approximately 1 mL EM with 0.01% MS-222 throughout the duration of the live imaging.

After larvae were immobilized, three baseline images were acquired in 5 min intervals. Then, 4 – 5 volumes of EM containing 0.01% MS-222 (1 mL each) with or without cisplatin (Abcam; ab141398) were exchanged around the mounted fish to initiate the exposure protocol. After 2 hours of exposure, fish were rinsed with 4 – 5 volumes of EM containing 0.01% MS-222 (1 mL each), maintained under sedation for an additional 2 – 4 hours depending on the specific experiment, and then euthanized.

For experiments with *mitoTimer* larvae, imaging was obtained before and after the cisplatin exposure protocol. For time-lapse experiments, images were acquired every 10 minutes throughout the 4-hour exposure protocol.

### Experiments involving Rho-Mito

Rho-Mito is a vital dye that localizes within mitochondria and fluoresces in the presence of cisplatin ^42^. Larvae at 5 – 7 dpf expressing either GCaMP3 in mitochondria (*Tg(myo6b:mitoGCaMP3))* or GFP in supporting cells (*Tg(TNKS1bp1:EGFP))* were incubated in EM with or without 1 mM cisplatin for either 30 or 60 min, depending on the experiment. This was followed by three rinses in EM. They were then submerged in EM containing Rho-Mito 3 µM for 15 minutes. Next, the larvae were rinsed twice in EM and then sedated and mounted for live imaging as previously described. Larvae expressing *mitoGCaMP3* were imaged at baseline and after 4 additional hours of recovery in EM to observe subsequent hair cell death. Larvae expressing GFP in supporting cells were only imaged at their specified time point to evaluate the dynamics of cisplatin accumulation within mitochondria.

The original description of Rho-Mito suggests pre-labeling cultured cells *in vitro* and then applying cisplatin ^42^. However, when optimizing our experimental approach, we observed Rho-Mito signal was barely detectable in our preloaded hair cells after completion of our cisplatin exposure protocol, which may be due to efflux of Rho-Mito from the zebrafish hair cells through passive and/or active transport. Since Rho-Mito fluoresces rapidly in the presence of cisplatin, we modified the protocol by loading hair cells with Rho-Mito immediately after they had been exposed to cisplatin just prior to imaging

### Image acquisition and analysis for live imaging

Z-stack images (2 µm step size; 30 steps per Z-stack) of neuromasts L2 – L4 were obtained with an ORCA-Flash 5.0 V3 camera (Hamamatsu) using an X-Light V2TP spinning disk confocal and a 63x/0.9N.A. water immersion objective on a Leica DM6 Fixed Stage microscope. Green fluorescence from *mitoGCaMP3-* and *cytoGCaMP3-*labeled hair cells and GFP-labeled supporting cells were imaged using a 470 nm wavelength laser at 20% power with a 50 ms/frame exposure time. Red fluorescence from hair cells labeled with Rho-Mito were imaged using a 555 nm wavelength laser at 20% power with a 50 ms/frame exposure time. In *mitoTimer*-labeled hair cells, green and red fluorescence were measured simultaneously using a Cairn OptoSplit II Bypass Image Splitter (Cairn Research Ltd). Multi-position image acquisition was accomplished using a Motorized ZDeck Quick-Adjust Platform System (Prior Scientific) directed by MetaMorph software (Molecular Devices).

Digital images of neuromasts were processed and analyzed using ImageJ software ^67^. Evaluation of changes in fluorescent intensity of genetically encoded biosensors was performed as previously described with a few modifications ^68^. In brief, maximum intensity projections of each z-stack were generated, then background subtraction was performed using a rolling ball radius of 100 pixels. A circular region of interest (ROI) with a diameter of 4 µm was placed around each hair cell according to its baseline position. Average fluorescence intensities within the ROI were then measured and expressed relative to the median baseline intensity of control hair cells. For time-lapse experiments, Stackreg and Turboreg ImageJ plugins were used for recursive alignment of hair cells that drifted over the acquisition time ^69^. Hair cells were characterized as living or dying based on their fragmentation and neuromast clearance (as described in ^37^). Briefly, hair cells were enumerated prior to cisplatin exposure. Intact hair cells that were still part of the neuromast structure at completion of the cisplatin exposure protocol were considered alive. Hair cells that underwent clearance from the neuromast, defined as disintegration of their cell membrane followed by ejection from the neuromast, were considered dead. Data were excluded at time points when fragments of dying hair cells drifted into neighboring ROIs of living hair cells.

### Microflume protocol

Experiments with the microflume were adapted from methods that used an orbital shaker to stimulate the lateral-line organ ^23^. Briefly, larvae at 5 dpf were transferred into a microflume apparatus that produces a consistent flow stimulus of 5 mm/sec (∼1 larval length/sec) ^70^. Larvae were exposed to this flow stimulus for 24 hours in EM. Following exposure to microflume stimulation, larvae were immobilized and baseline images were acquired. Then they were exposed to 250 µM cisplatin for 2 hours and imaged after an additional 4 hours of recovery in EM with 0.01% tricaine. Imaging processing and analysis were performed as in the other live imaging experiments with the following modifications. Images were acquired with an exposure time of 100 ms/frame instead of 50 ms/frame. The ratio of *mitoTimer* red:green fluorescence for each hair cell was calculated and normalized to the median value of control hair cells. Hair cell counts per neuromast were expressed as a percentage of remaining intact hair cells after cisplatin exposure compared to baseline.

### Whole-mount immunohistochemistry, imaging, and analysis of caspase-3 experiments

Cisplatin exposure and fixation with activated caspase-3 antibodies were performed as previously described with a few modifications ^18^. Briefly, groups of wildtype larvae (AB*) at 6 dpf were incubated for 2 hours in 30 mL EM with or without 1 mM cisplatin (Abcam) at 28°C. Larvae were then rinsed and allowed to recover for an additional 2 hours in 30 mL EM. At every 30-minute increment starting immediately prior to treatment initiation, groups of control and cisplatin-exposed larvae were euthanized by rapid chilling and fixed overnight in 4% paraformaldehyde in 0.1 M phosphate-buffered solution (PBS pH = 7.4) at 4°C. The next day, fixed specimens were rinsed, blocked, and incubated in primary antibodies (1:500 HCS-1 mouse monoclonal, DSHB, University of Iowa; 1:400 anti-cleaved caspase-3 rabbit polyclonal, Cell Signaling Technology). After overnight incubation in primary, specimens were rinsed and incubated in DAPI (5 µg/mL) and secondary antibodies (1:500 anti-mouse IgG conjugated to Alexa Fluor 488 and 1:500 anti-rabbit IgG conjugated to Alexa Fluor 555) at room temperature in darkness. Specimens were rinsed in PBS prior to being mounted on glass slides with elvanol (13 % w/v polyvinyl alcohol, 33 % w/v glycerol, 1 % w/v DABCO (1,4 diazobicylo[2,2,2] octane) in 0.2 M Tris, pH 8.5).

The posterior lateral-line neuromasts (L3 – L5) of fixed and labeled zebrafish were imaged as in the prior live imaging experiments except that images were acquired with an exposure time of 100 ms/frame and a step size of 1 µm. DAPI-stained hair cells were acquired with a 405 nm wavelength laser. The percentage of hair cells with cleaved caspase-3 were determined for each neuromast and then averaged across neuromasts present within the same fish to account for fish-to-fish variability.

### Statistics

Standard descriptive statistics were used to describe experimental results. The two groups were determined using the unpaired Student’s *t*-test or Mann-Whitney *U* test, as appropriate. Comparison of multiple groups was performed by one-way ANOVA or Kruskal-Wallis test depending on data normality with appropriate *post hoc* tests as needed. A mixed model with Bonferroni *post hoc* analysis was performed on the cleaved caspase-3 experiments. All statistical analyses were conducted using Prism 9 (GraphPad Software Inc).

### Study approval

All experiments on zebrafish were performed with the approval of the Institutional

Animal Care and Use Committee of Washington University School of Medicine in St.

Louis (Protocol number: 20-0158) and in accordance with NIH guidelines for use of zebrafish.

### Data availability

Underlying graphed data and reported means can be found in the Supporting Data Values file. STL files of microflume components are available in the Open Science Framework repository, https://osf.io/rvyfz/.

## Supporting information

Movie 1 - mitoGCaMP3

Movie 2 - cytoGCaMP3

Supplemental Datasharing

## Author contributions

DSL (conceptualization, investigation, methodology, formal analysis, visualization, writing – original draft, writing – review and editing, funding acquisition), AS (investigation), JZ (resources, methodology, writing – review and editing), WHA (resources, methodology, writing – review and editing), MW (conceptualization, supervision, writing – review and editing, funding acquisition), LS (conceptualization, supervision, project administration, formal analysis, visualization, writing – original draft, writing – review and editing, funding acquisition)

## Acknowledgements

Research reported in this publication was supported by a Large-Scale Interdisciplinary Research Initiative from the Children’s Discovery Institute, St. Louis Children’s Hospital (LS and MW; Grant # MC-LI-2018-762), the American Society of Pediatric Otolaryngology CORE grant (DL.; # 930395), and the National Institute of Deafness and Other Communication Disorders (NIDCD) within the National Institutes of Health (NIH), through the “Development of Clinician/Researchers in Academic ENT” training grant, award number T32DC000022 (DL). The content is solely the responsibility of the authors and does not necessarily represent the official view of the funding sources. We further acknowledge Emily L. Bell for her technical support in the live *in vivo* imaging experiments.

